# Amniotic fluid stem cell extracellular vesicles promote fetal lung branching and cell differentiation in experimental congenital diaphragmatic hernia

**DOI:** 10.1101/2022.01.10.475632

**Authors:** Kasra Khalaj, Rebeca Lopes Figueira, Lina Antounians, Sree Gandhi, Matthew Wales, Louise Montalva, George Biouss, Augusto Zani

## Abstract

Pulmonary hypoplasia secondary to congenital diaphragmatic hernia (CDH) is characterized by impaired branching morphogenesis and differentiation. We have previously demonstrated that administration of extracellular vesicles derived from rat amniotic fluid stem cells (AFSC-EVs) rescues development of hypoplastic lungs at the pseudoglandular and alveolar stages in rodent models of CDH. Herein, we tested whether AFSC-EVs exert their regenerative effects at the canalicular and saccular stages, as these are translationally relevant for clinical intervention. To induce fetal pulmonary hypoplasia, we gavaged rat dams with nitrofen at embryonic day 9.5 and demonstrated that nitrofen-exposed lungs had impaired branching morphogenesis, dysregulated signaling pathways relevant to lung development (FGF10/FGFR2, ROBO/SLIT, Ephrin, Neuropilin 1, β-catenin) and impaired epithelial and mesenchymal cell marker expression at both stages. AFSC-EVs administered to nitrofen-exposed lung explants rescued airspace density and increased the expression levels of key factors responsible for branching morphogenesis. Moreover, AFSC-EVs rescued the expression of alveolar type 1 and 2 cell markers at both canalicular and saccular stages, and restored markers of club, ciliated epithelial, and pulmonary neuroendocrine cells at the saccular stage. AFSC-EV treated lungs also had restored markers of lipofibroblasts and PDGFRA+ cells to control levels at both stages. EV tracking showed uptake of AFSC-EV RNA cargo throughout the fetal lung and an mRNA-miRNA network analysis identified that several miRNAs responsible for regulating lung development processes were contained in the AFSC-EV cargo. These findings suggest that AFSC-EV based therapies hold potential for restoring fetal lung growth and maturation in babies with pulmonary hypoplasia secondary to CDH.

**Graphical abstract:** Babies with congenital diaphragmatic hernia have hypoplastic lungs characterized by impaired branching morphogenesis and undifferentiated epithelium and mesenchyme. The authors demonstrate that amniotic fluid stem cell extracellular vesicles (AFSC-EVs) administered to rat hypoplastic fetal lungs restore branching and exert regenerative effects on epithelial and mesenchymal cells, partly through miRNA cargo transfer. AFSC-EV beneficial effects were obtained at translationally relevant timepoints.

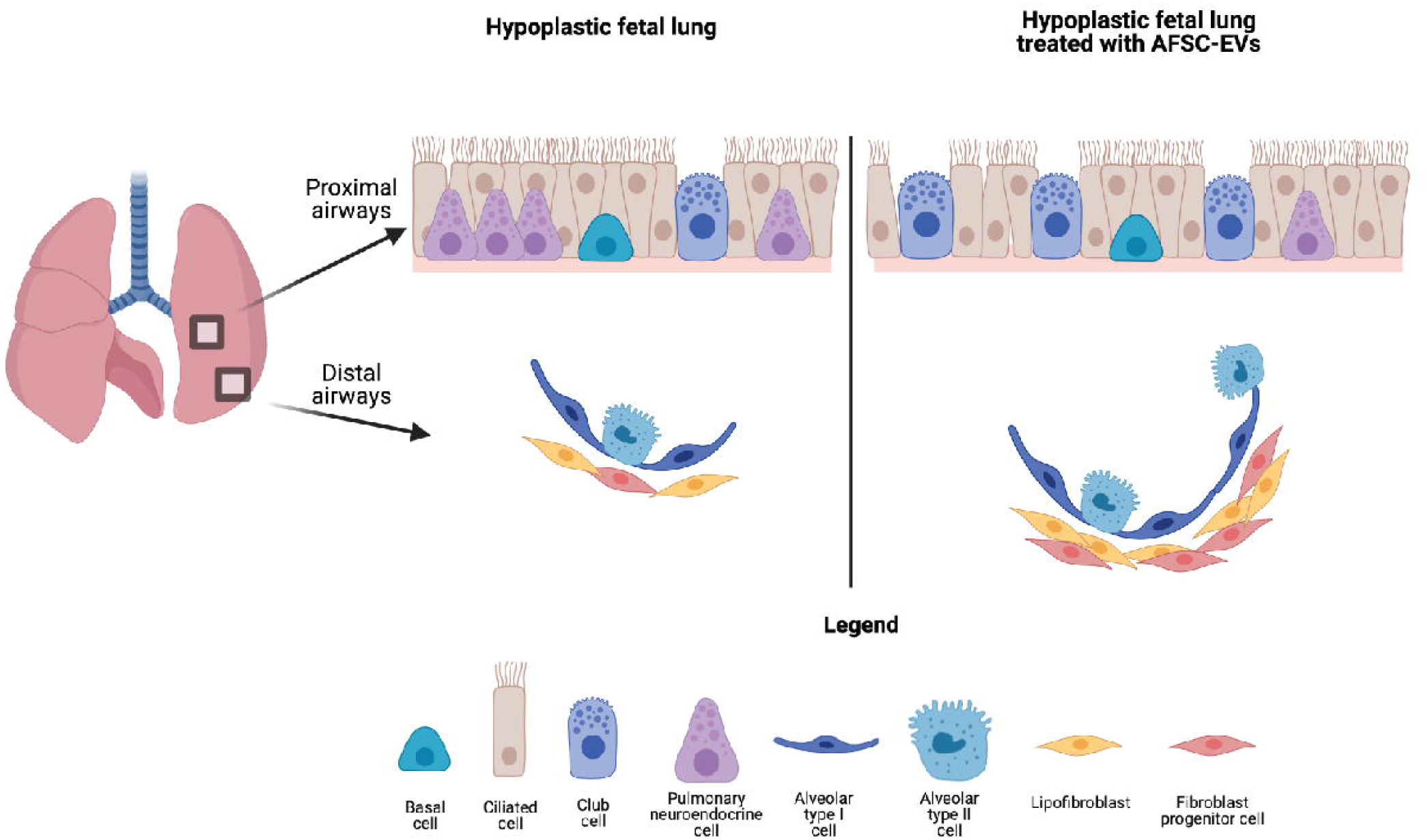

## Introduction

Fetal lung underdevelopment, also known as pulmonary hypoplasia, is primarily characterized by decreased lung growth and maturation [1]. The most common birth defect found in babies with pulmonary hypoplasia is congenital diaphragmatic hernia (CDH). CDH is characterized by incomplete closure of the fetal diaphragm and herniation of abdominal organs into the chest. At present, the mortality of CDH is 20-30% [2,3], and around 60% of survivors have long-term morbidity that may persist into adulthood [4]. Pulmonary hypoplasia is a primary cause of these poor outcomes, and it persists even after surgical repair of the diaphragm and repositioning of the herniated organs back into the abdomen [5]. The poor outcomes observed in infants and children with CDH have mobilized researchers to find an antenatal treatment that can promote normal lung development in fetuses with pulmonary hypoplasia secondary to CDH [6].

We have recently reported that administration of extracellular vesicles derived from rat amniotic fluid stem cells (AFSC-EVs) rescues normal development in rodent fetal models of pulmonary hypoplasia [7]. Specifically, we demonstrated that AFSC-EVs increase branching morphogenesis and promote epithelial cell differentiation in fetal rat lung explants harvested at the pseudoglandular stage [7]. Moreover, AFSC-EVs administered intra-tracheally in an *in vivo* rabbit fetal model of CDH increased the number of alveoli and promoted lipofibroblast differentiation at the alveolar stage of lung development [7]. The effects of AFSC-EVs on rat fetal lung growth and maturation were mainly mediated by their RNA cargo, as enzymatically digesting AFSC-EV RNA ablated their beneficial effects in multiple models [7]. Of the small RNA species contained in AFSC-EV cargo, miRNAs were the species most likely to mediate the regenerative effects observed on lung development [7].

When we previously investigated the biological effects of AFSC-EVs on fetal hypoplastic lungs, we reported that Fibroblast growth factor 10 (FGF10) expression was rescued to normal levels [7]. In that study, we had focused on the FGF10 pathway, as it plays a critical role in lung branching morphogenesis, epithelial proliferation, and lineage commitment [8]. However, fetal lung development is a highly dynamic process that requires multiple pathways to coordinate branching morphogenesis and lung cell differentiation [9]. Building on recent studies on the transcriptomic analysis and mapping of the rodent fetal lung [10,11], herein we investigated multiple signaling pathways that are known or unknown to be dysregulated in experimental CDH. The findings reported in the present study expand on our knowledge of pulmonary hypoplasia and further elaborate on the impact of AFSC-EVs on regulatory pathways that are key for lung development. Moreover, we further delineated the effects of AFSC-EV therapy on hypoplastic lungs at the canalicular and saccular stages. As CDH is typically diagnosed at the anatomy scan around 20 weeks of gestation, these stages are translationally relevant and provide an opportunity for designing clinical interventional studies. Lastly, the present study further characterized the AFSC-EV effects on epithelial and mesenchymal cell differentiation, specifically interrogating some of the most relevant cell types during the canalicular and saccular stages of lung development.

## Materials and Methods

### Extracellular vesicle isolation and characterization

Extracellular vesicles were derived from c-kit+ rat amniotic fluid stem cells (AFSCs) using our established protocol [12]. Briefly, AFSCs were grown to 70% confluency and the supernatant was collected and subjected to differential centrifugation (300 g, 1,200 g, and 100,000 g), as described [12]. Following the International Society for Extracellular Vesicles guidelines [13], EVs were characterized for size (nanoparticle tracking analysis), morphology (transmission electron microscopy), and expression of canonical EV-related protein markers (Western blot analysis), as described [7,12].

### Fluorescence labelling of AFSC-EVs and in vitro tracking

AFSC-EVs were fluorescently labelled using ExoGlow-EV labeling kit (SBI System Biosciences), using the manufacturer’s protocol. The fluorescently labelled AFSC-EVs were added to E17.5 and E20.5 fetal rat explant cultures and co-incubated with 4′,6-diamidino-2-phenylindole (DAPI) for 3 hrs prior to imaging, using a 2-photon Leica SP8 confocal microscope (Leica).

### Experimental models of pulmonary hypoplasia

#### Ex vivo

In fetal rats, CDH and pulmonary hypoplasia were induced via the administration of nitrofen to rat dams (100 mg) by oral gavage on embryonic day (E) 9.5. Dams were euthanized and fetal lungs were micro-dissected at the canalicular (E17.5) and saccular (E20.5) stages of lung development. Lungs were grown as explants on nanopore membranes (Whatman, Thermofisher Scientific), and incubated for 72 h in culture medium alone or rat AFSC-Evs (0.5% v/v). Lungs from fetuses of dams that received olive oil (no nitrofen) at E9.5 served as control. All explant specimens were snap frozen in liquid nitrogen and stored at -80 °C for long-term storage or fixed and processed for histology using our established protocol [7].

#### In vitro

Primary fibroblasts were derived from fetal rat lungs dissected at E18.5 using an established protocol [14]. Briefly, single cell suspensions were obtained from pooled lungs of control and nitrofen-exposed rat fetuses by trypsinization (0.25% trypsin; Gibco). Fibroblasts were isolated as described [7,14,15], grown for 7 days and then treated with culture medium alone or rat AFSC-EVs (0.5% v/v) for 24 h. Upon harvest, fibroblast gene expression analysis was conducted.

### Branching morphogenesis evaluation

To assess branching, rat fetal lung explants were processed for histology (hematoxylin/eosin staining) as described [7]. Explants were evaluated for airspace density using the radial airspace count (RAC) and parenchymal to airspace ratio using the mean linear intercept (MLI), as recommended by the American Thoracic Society [16].

### mRNA quantification with qRT-PCR

RNA extraction was conducted using Trizol and cDNA synthesis was performed using the VILO superscript RT III kit (Life Technologies), as described [7]. Gene expression was conducted using the 2^-deltadeltaCT method using ViiA qPCR system (ABI) following MIQE guidelines [17]. Primers are listed in Table 1.

**Table 1.**
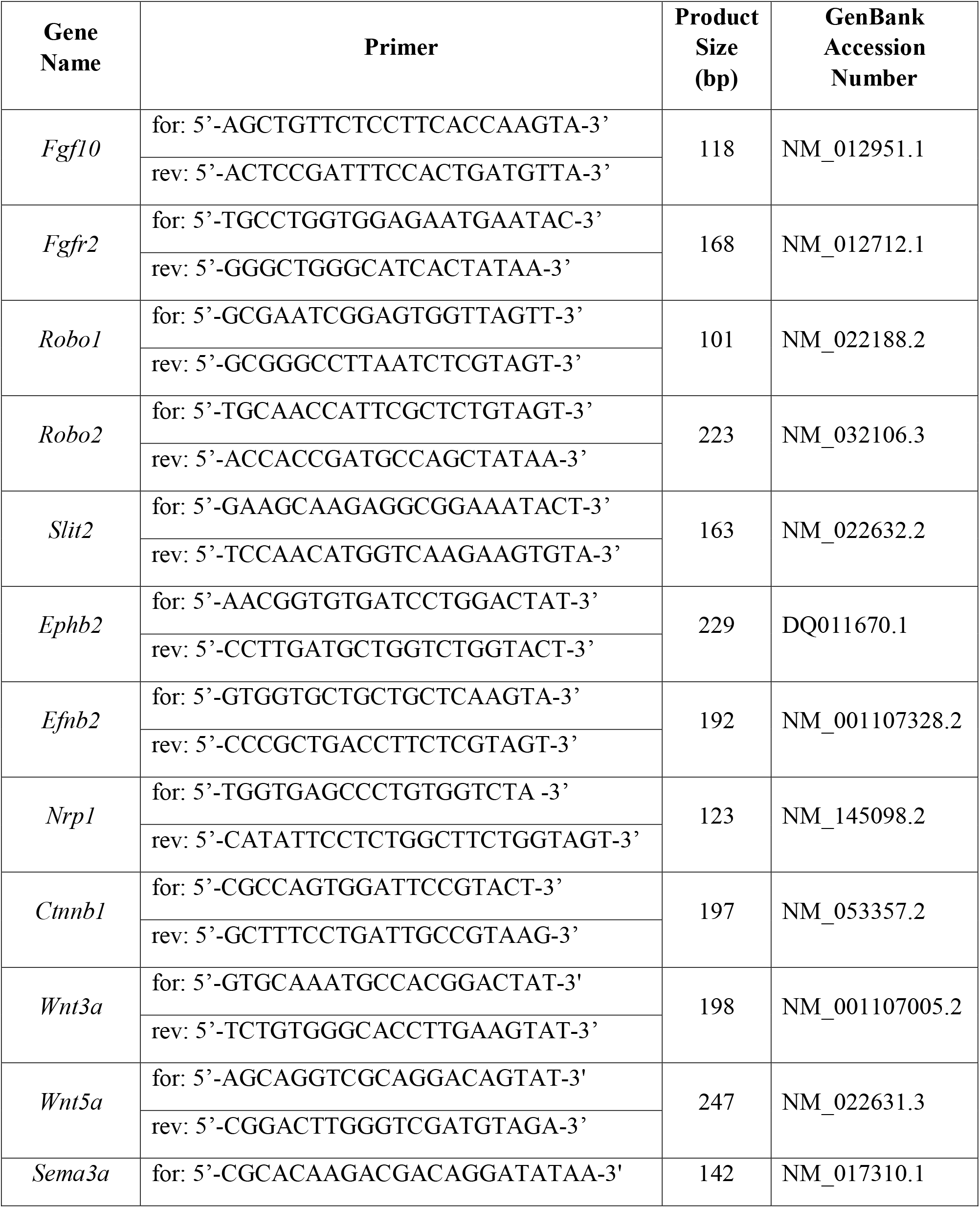
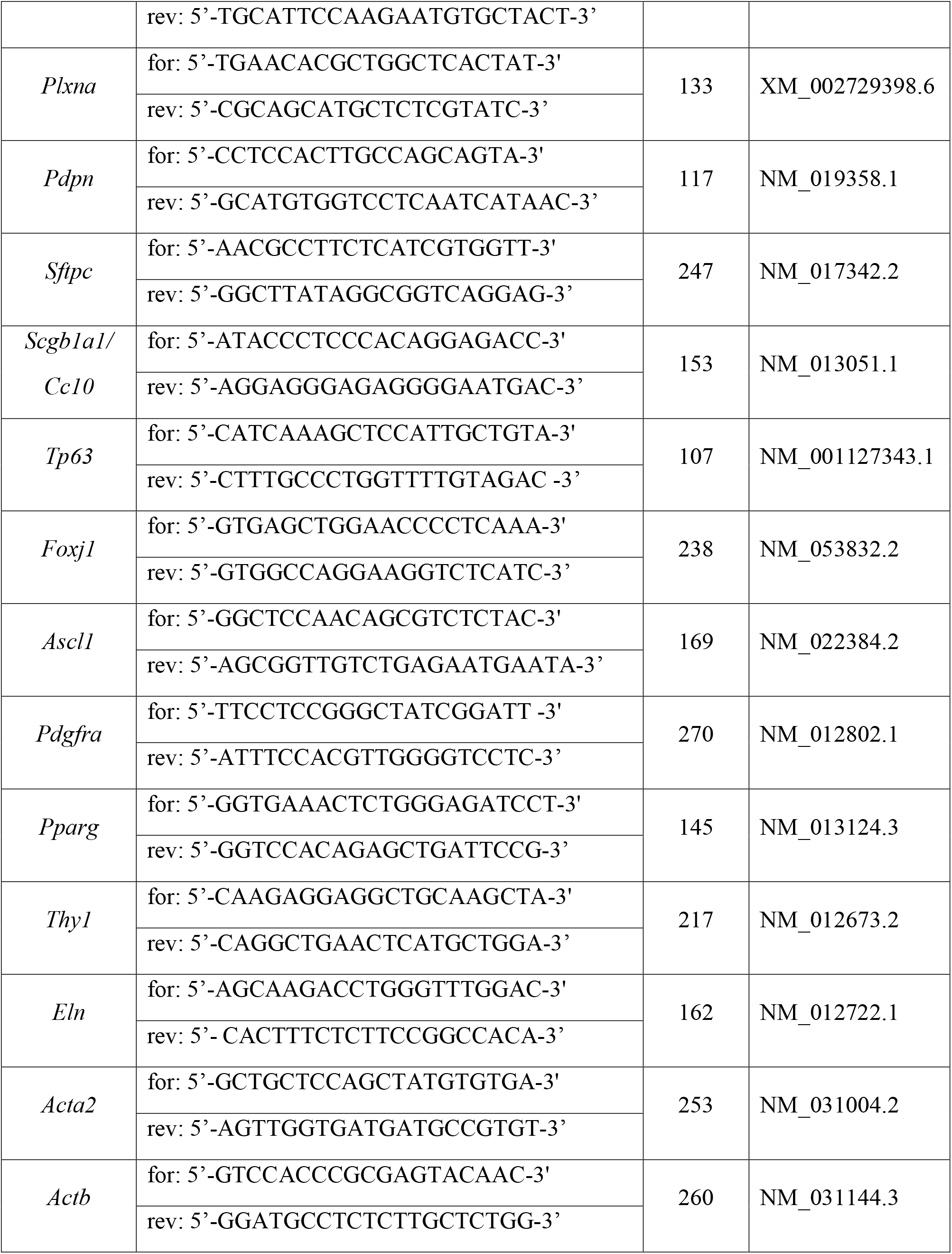
mRNAs assessed by Real-Time PCR.

### Protein extraction and Western blotting

Protein extraction and Western blotting were performed, as previously described [7]. Briefly, cells or tissues were subjected to tissue extraction buffer (ThermoFisher Scientific) containing protease and phosphatase inhibitor cocktail (Roche) and disrupted by sonication. The protein content was quantified using a bicinchoninic acid assay (ThermoFisher Scientific), normalized to 10 μg, and loaded on 4-12% NuPage MES mini-blot pre-cast gels (ThermoFisher Scientific). Transfer was performed using the iBlot2 system (ThermoFisher Scientific) and imaged using an Odyssey FC digital acquisition system (LI-COR Biosciences). Antibodies and concentrations are specified in Table 2.

**Table 2.** Antibodies utilized for imaging and Western blotting.

| **Antibody** | **Antibody host** | **Manufacturer/Catalogue #** | **Application** | **Dilution Factor** |
| --- | --- | --- | --- | --- |
| PDPN | Rabbit | ThermoFisher/P5374 | WB, IF | 1:200 (IF), 1:1000 (WB) |
| EPCAM | Rabbit | Abcam/ab213500 | IF | 1:100 |
| SPC | Rabbit | Abcam/ab90716 | WB, IF | 1:400 (IF), 1:1000 (WB) |
| ECAD | Mouse | Abcam/ab231303 | IF | 1:100 |
| CC10 | Mouse | Santa-Cruz/Sc-3659921 | WB, IF | 1:100 (IF), 1:1000 (WB) |
| FOXJ1 | Mouse | Santa-Cruz/Sc-53139 | WB, IF | 1:50 (IF), 1:500 (WB) |
| P63 | Rabbit | Abcam/ab124762 | WB, IF | 1:50 (IF), 1:500 (WB) |
| ASCL1 | Rabbit | Invitrogen/14579482 | WB,IF | 1:250 (IF), 1:1000 (WB) |
| PPARG | Rabbit | ThermoFisher/PA3 821A | WB, IF | 1:500 (IF), 1:1000 (WB) |
| PDGFRA | Goat | R&D Systems/AF1062 | WB, IF | 1:500 (IF), 1:1000 (WB) |
| IgG Alexafluor-488 | Rabbit | CST/4412S | IF | 1:1000 |
| IgG Alexafluor-555 | Mouse | CST/4409 | IF | 1:1000 |
| IgG AlexaFluor-488 | Goat | Invitrogen/A11055 | IF | 1:1000 |
| IgG HRP | Mouse | CST/7076S | WB | 1:1000 |
| IgG HRP | Rabbit | CST/7074 | WB | 1:1000 |
| IgG HRP | Goat | Invitrogen/31402 | WB | 1:1000 |

CST: Cell signaling technology
WB: Western blot
IF: Immunofluorescence

### Immunofluorescence using confocal microscopy

Immunofluorescence was conducted using our established protocol [7]. Paraffin-embedded sections were stained using the following antibodies: Podoplanin (PDPN) for alveolar type 1 (AT1) cells, Surfactant protein C (SPC) for alveolar type 2 (AT2) cells, both co-stained with Epithelial adherens junction adherin (ECAD); Club-cell-specific 10 kD protein (CC10) for club cells; Tumor protein 63 (P63) for basal cells; Forkhead box protein J1 (FOXJ1) for ciliated epithelial cells; Achaete-scute homolog 1 (ASCL1) for pulmonary neuroendocrine cells; Platelet-derived growth factor receptor A (PDGFRA) and Peroxisome proliferator-activated receptor gamma (PPARG) for mesenchymal cells. Antibodies and concentrations are provided in Table 2.

## Results

### Administration of AFSC-EVs rescues branching morphogenesis in fetal hypoplastic lungs during the canalicular and saccular stages of lung development

We first confirmed that nitrofen-exposed lungs harvested at both canalicular and saccular stages of lung development had impaired branching morphogenesis, as evidenced by fewer airspaces measured with RAC and a lower airspace to parenchyma ratio measured by MLI (**Figure 1A-B**). Administration of AFSC-EVs to nitrofen-exposed lung explants rescued branching morphogenesis at both developmental stages. When we investigated signaling pathways that are known to be responsible for branching morphogenesis [8], we found that nitrofen-exposed lungs harvested at either E17.5 or E20.5 had downregulated expression of *Fgf10*, Roundabout guidance receptor 2 (*Robo2*), and Slit guidance ligand 2 (*Slit2*) factors, as well as upregulated expression of Roundabout guidance receptor 1 (*Robo1*) (**Figure 1C-D**). Moreover, fetal lungs harvested at E20.5 had higher levels of Fibroblast growth factor receptor 2 (*Fgfr2*) and lower levels of Ephrin B2 (*Efnb2*), Ephrin type-B receptor 2 (*Ephb2*), Neuropilin 1 (*Nrp1*), and β-catenin 1 (*Cttnb1*) compared to control (**Figure 1D**). Interestingly, nitrofen-exposed lungs did not have any dysregulation of other factors that are also involved in branching morphogenesis [8], such as Wnt family member 3a and 5a (*Wnt3a* and *Wnt5a*), Semaphorin 3a (*Sema3a*), and Plexin A (*Plxna*) at any of the timepoints (**Supplementary Figure 1**). Following AFSC-EV administration, nitrofen-exposed lungs harvested at E17.5 or E20.5 had upregulated levels of *Fgf10, Robo2*, and *Slit2* and downregulated levels of *Fgfr2* and *Robo1*. Moreover, AFSC-EV treated hypoplastic lungs harvested at E20.5 had upregulated levels of *Efnb2, Ephb2*, and *Nrp1* (**Figure 1C-D**).

**Figure 1.**
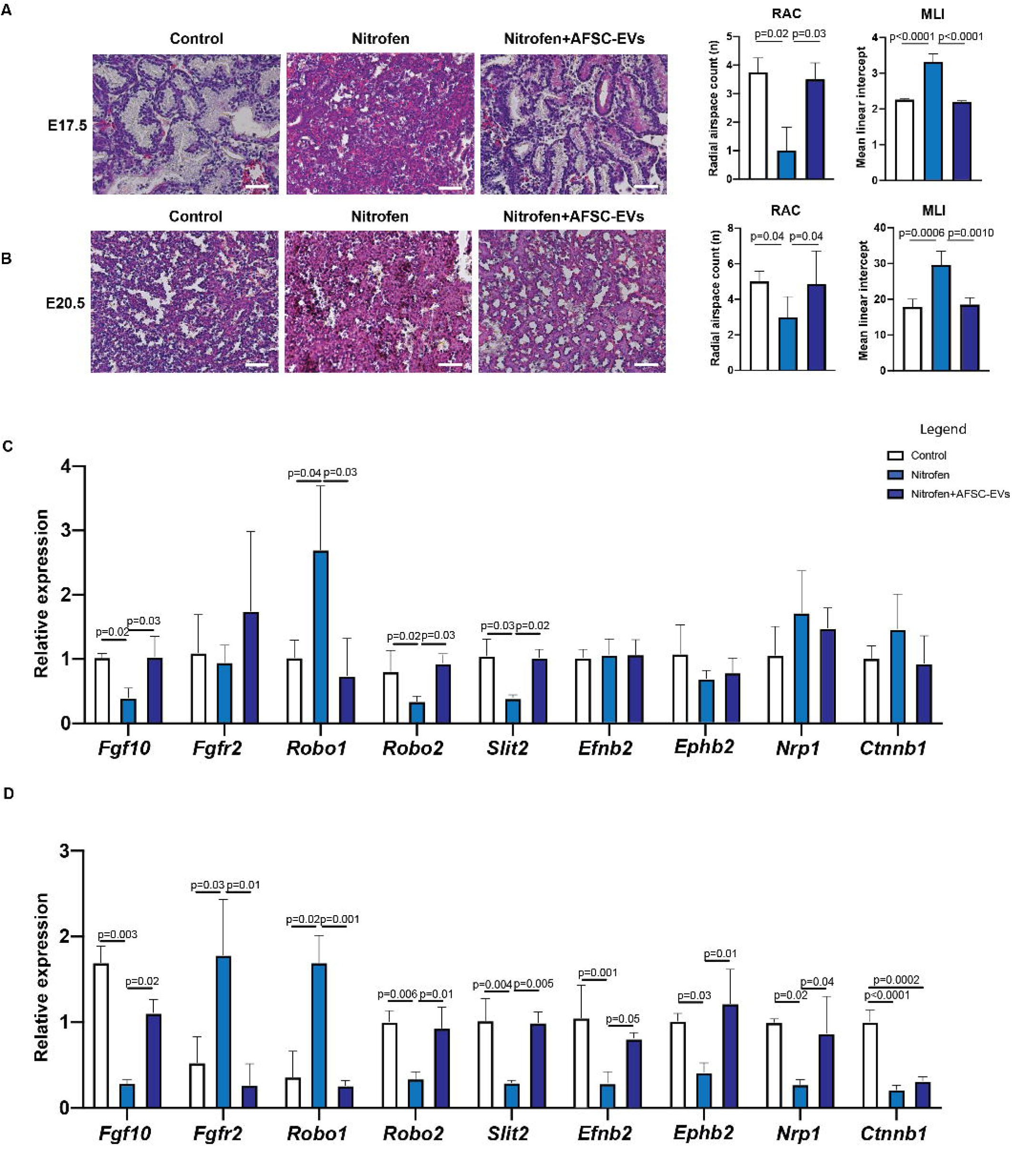
Administration of AFSC-EVs rescues branching morphogenesis in fetal hypoplastic lungs during the canalicular and saccular stages of development. Representative images of hematoxylin and eosin stains and quantification of airspace density using the radial airspace count (RAC) and parenchymal to airspace ratio using the mean linear intercept (MLI), in A, canalicular and B, saccular stages of lung development. Scale bar = 50 μm. Gene expression levels of factors involved in branching morphogenesis in control and nitrofen-exposed untreated and AFSC-EV treated lung explants at the C, canalicular and D, saccular stages of lung development. Data were analyzed using Kruskal-Wallis test for E17.5 MLI and RAC and E20.5 RAC (A,B) and one-way ANOVA using Tukey’s multiple comparison test for all other analyses (A-D). Data are presented as mean ± SD (n= at least 4 biological replicates per group for RAC and MLI; n= 3 biological replicates per group for gene expression analysis). Only significant differences (p<0.05) are reported in the graphs.

### AFSC-EV administration promotes differentiation of epithelial lung cells at canalicular and saccular stages of lung development

Nitrofen-exposed lungs have been reported to be immature, in part due to the altered differentiation of the proximal and distal airway epithelium. At the canalicular stage, we observed that nitrofen exposure decreased the expression of markers for AT1 (PDPN) and AT2 (SPC) cells but had no effect on markers of club cells (CC10), basal cells (P63), ciliated epithelial cells (FOXJ1), and pulmonary neuroendocrine cells (ASCL1) (**Figure 2A-C, Supplementary Figure 2**). Administration of AFSC-EVs to nitrofen-exposed lungs harvested at the canalicular stage rescued the expression levels of AT1 and AT2 cell markers back to control.

**Figure 2.**
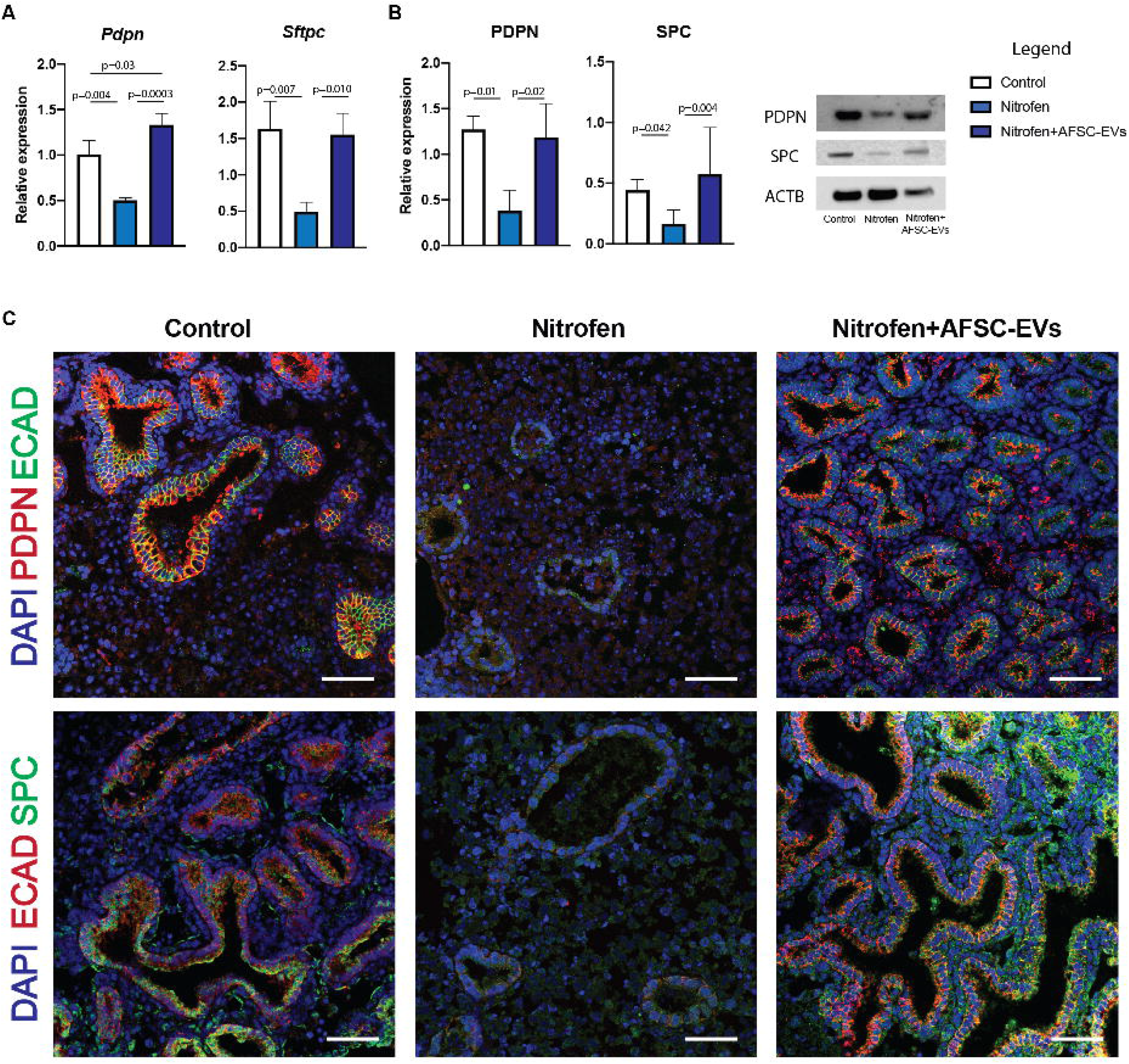
AFSC-EV administration promotes differentiation of alveolar lung epithelial cells at the canalicular stage of fetal lung development. Markers of AT1 (PDPN) and AT2 (SPC) were analyzed at the A, gene (RT-qPCR), and B, protein (Western blot) expression level. ACTB was used as a loading control. C, Representative immunofluorescence images of PDPN (red) and SPC (green) co-localization with the epithelial marker E-cadherin (ECAD). DAPI was used as a nuclear stain. Scale bar = 50 μm. Data were analyzed using one-way ANOVA using Tukey’s multiple comparison test. Data are presented as mean ± SD (n= at least 3 biological replicates per group for gene and protein expression). Only significant differences (p<0.05) are reported in the graphs.

At the saccular stage, we observed a dysregulation of expression markers of proximal and distal epithelial cells (**Figure 3A-C**). Nitrofen-exposed lungs harvested at this stage and treated with AFSC-EVs had increased expression of AT1, AT2, club, and ciliated epithelial cells, and decreased expression of pulmonary neuroendocrine cells back to control (**Figure 3A-C**).

**Figure 3.**
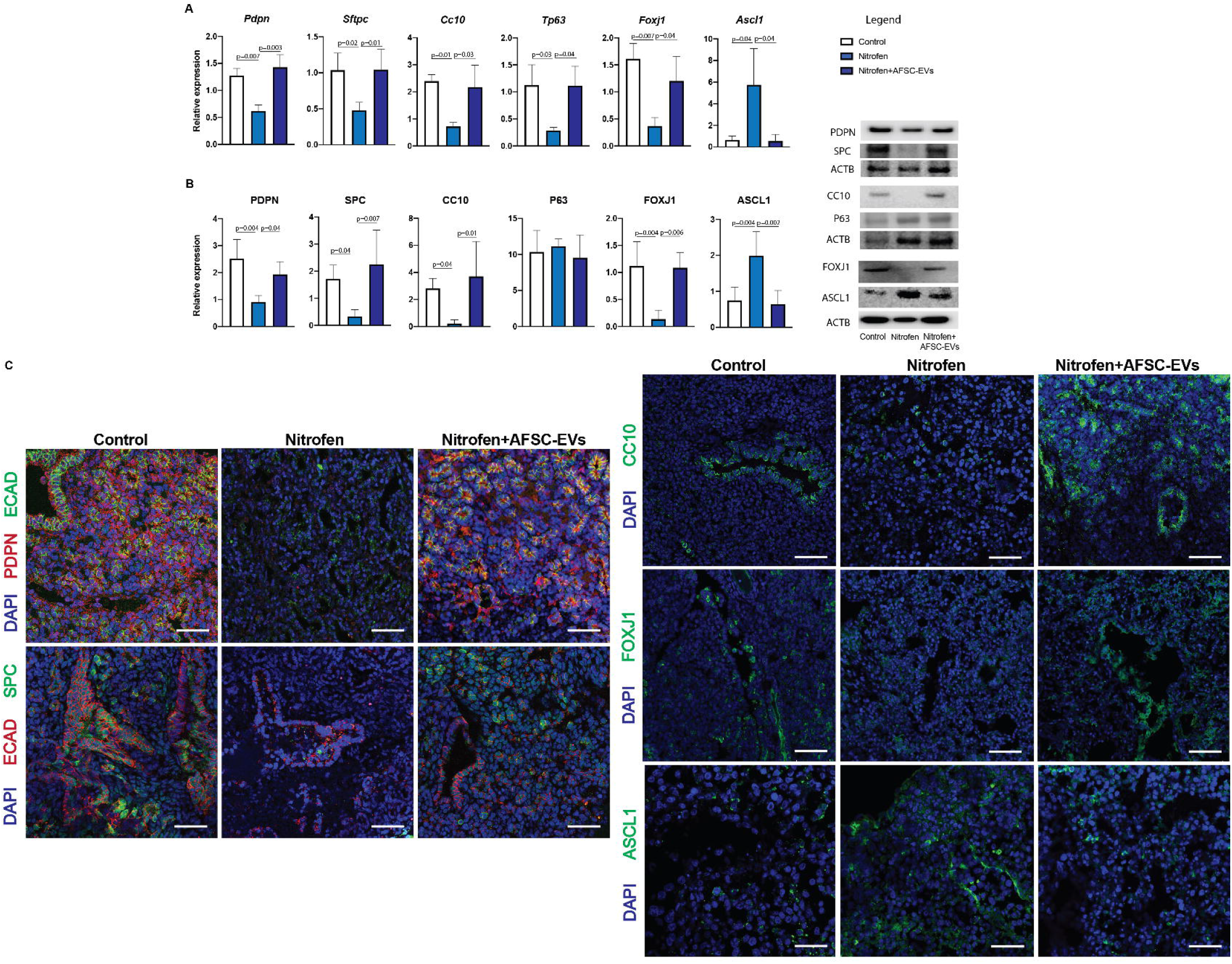
AFSC-EV administration promotes differentiation of alveolar and bronchial lung epithelial cells at the saccular stage of fetal lung development. Markers of AT1 (PDPN), AT2 (SPC), club (CC10), basal (P63), ciliated epithelial (FOXJ1) and pulmonary neuroendocrine (ASCL1) were analyzed at the A, gene (RT-qPCR), and B, protein (Western blot) expression level. ACTB was used as a loading control. C, Representative immunofluorescence images of PDPN (red) and SPC (green) co-localization with the epithelial marker ECAD and ASCL1, CC10, FOXJ1 (green) immunolocalization in all fetal lung groups. DAPI was used as a nuclear stain. Scale bar = 50 μm. Data were analyzed using one-way ANOVA using Tukey’s multiple comparison test. Data are presented as mean ± SD (n= at least 3 biological replicates per group for gene and protein expression). Only significant differences (p<0.05) are reported in the graphs.

### AFSC-EV administration has regenerative effects on the fetal lung mesenchyme

Several studies have reported that CDH hypoplastic lungs have undifferentiated mesenchyme [18,19], with altered expression of lipofibroblasts [20–23] and PDGFRA+ cells [24,25]. To study mesenchymal differentiation and lung fibroblast response to AFSC-EVs, we first derived primary fibroblasts from control and nitrofen-exposed lungs harvested at the canalicular stage. In primary cell culture, we observed a decreased expression of PPARG and Thy1, which are markers of lipofibroblasts [26]. Moreover, we found an increase in PDGFRA expression, which is a marker of progenitor cells that mainly give rise to myofibroblasts [27], but no difference in expression of Actin alpha 2 (*Acta2*), a well-established marker of myofibroblasts [18,19]. When we investigated the mesenchyme of fetal lung explants, we confirmed the decrease in lipofibroblast markers at both gene and protein expression levels in nitrofen-exposed fetal lungs harvested at canalicular (**Figure 4B-D**) and saccular stages (**Figure 4E-G**). The increase in PDGFRA+ cell expression noticed in primary fibroblasts derived from nitrofen-exposed lungs was also confirmed in the explant model at the canalicular stage (**Figure 4B-D**). Conversely, the protein expression of PDGFRA in fetal nitrofen-exposed lungs harvested at the saccular stage was decreased compared to control (**Figure 4F-G**). Administration of AFSC-EVs to nitrofen-exposed primary fibroblasts or lung explants increased the expression of lipofibroblast markers (**Figure 4A-G**). Similarly, nitrofen-exposed primary fibroblasts and fetal lungs treated with AFSC-EVs had a decrease in PDGFRA expression levels compared to nitrofen-injured untreated fibroblasts or lung explants (**Figure 4B-G**).

**Figure 4.**
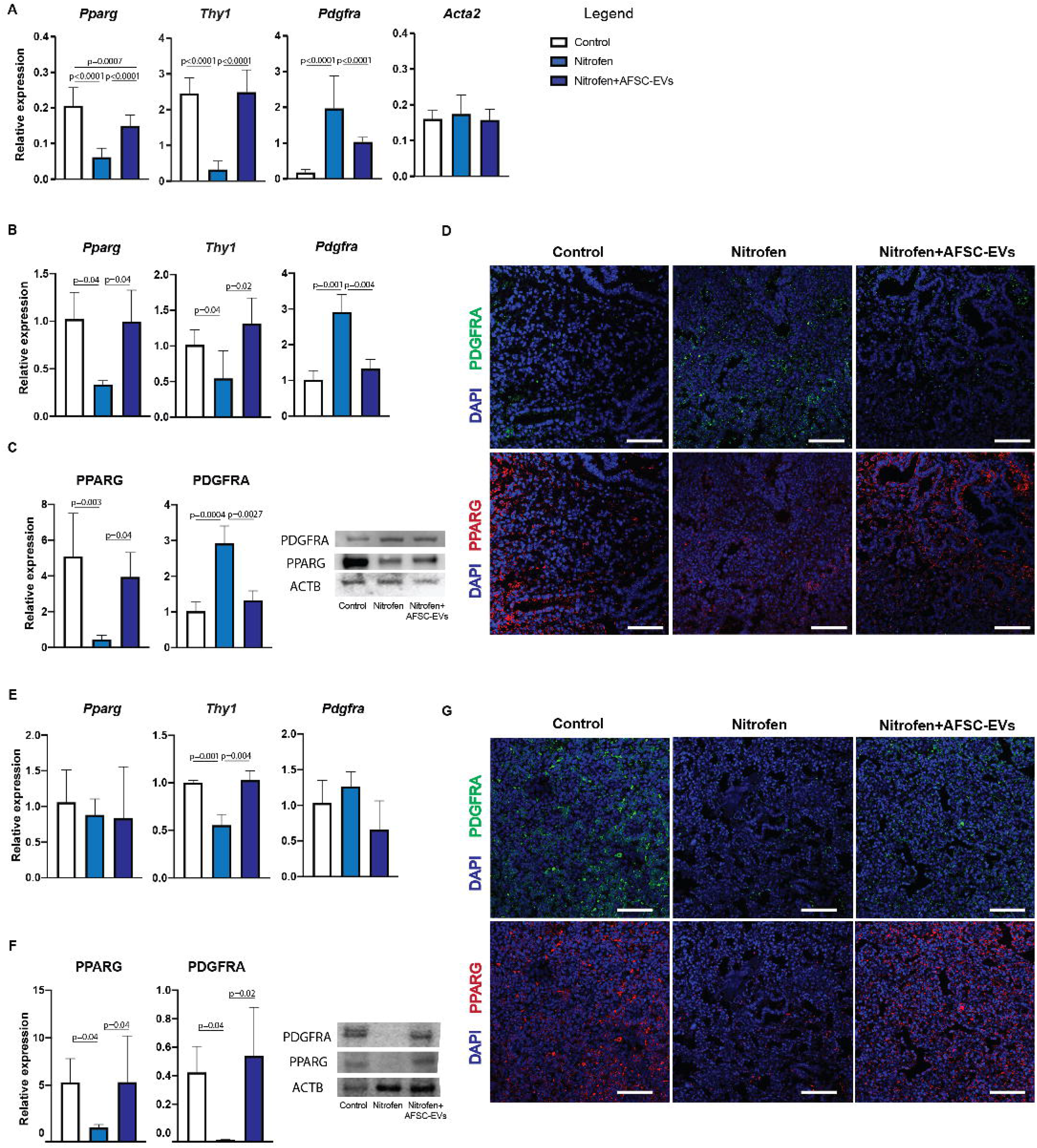
AFSC-EV administration has regenerative effects on the fetal lung mesenchyme. A, Mesenchymal markers for fibroblast (*Pdgfra*), lipofibroblast (*Pparg, Thy1*) and myofibroblast (*Eln, Acta2*) cells were analyzed in primary fibroblasts isolated from fetal lungs. Mesenchymal markers analyzed at the B, (RT-qPCR), and B, protein (Western blot) expression levels at the canalicular stage. ACTB was used as a loading control. D, Representative immunofluorescence images of the increased PDGFRA+ fibroblasts and diminished lipofibroblast cells at the canalicular stage. DAPI was used as a nuclear stain. Scale bar = 50 μm. These same markers were also analyzed at the E, gene and F, protein expression levels at the saccular stages. G, Representative immunofluorescence images of decreased PDGFRA+ fibroblast and lipofibroblast cells at the saccular stage. DAPI was used as a nuclear stain. Scale bar = 50 μm. Data were analyzed using one-way ANOVA using Tukey’s multiple comparison test. Data are presented as mean ± SD (n= at least 3 biological replicates per group for gene and protein expression). Only significant differences (p<0.05) are reported in the graphs.

### AFSC-EV mechanism of action is partly modulated by the release of RNA species

As we have previously demonstrated that AFSC-EV main mechanism of action towards fetal lung regeneration at E14.5 is through the release of the RNA cargo [7], in this study we tracked the RNA cargo of AFSC-EVs that were administered to nitrofen-exposed lungs harvested at E17.5 and E20.5. At both stages of lung development, we observed that AFSC-EVs entered the lung tissue, as shown by the presence of their RNA cargo throughout the lung parenchyma (**Figure 5A-B, Supplementary online Video 1**). In our previous small RNA sequencing studies [7], we identified some miRNAs that were in part responsible for the effects of AFSC-EVs on branching morphogenesis and fetal lung epithelial differentiation. In the present study, we linked the genes responsible for branching morphogenesis and cell differentiation that were downregulated following AFSC-EV treatment (*Robo1, Fgfr2, Ascl1*, and *Pdgfra*; **Figures 1,3,4**) with miRNAs that were found to be in the AFSC-EV cargo by the small RNA sequencing analysis conducted in our previous study (**Figure 5C**). Using miRWalk, we found 91 miRNAs that are predicted to downregulate the selected 4 genes, including 28 that are known to be involved in lung development (**Figure 5C**). Of the 91 miRNAs, we identified 11 miRNAs that co-regulated more than one gene.

**Figure 5.**
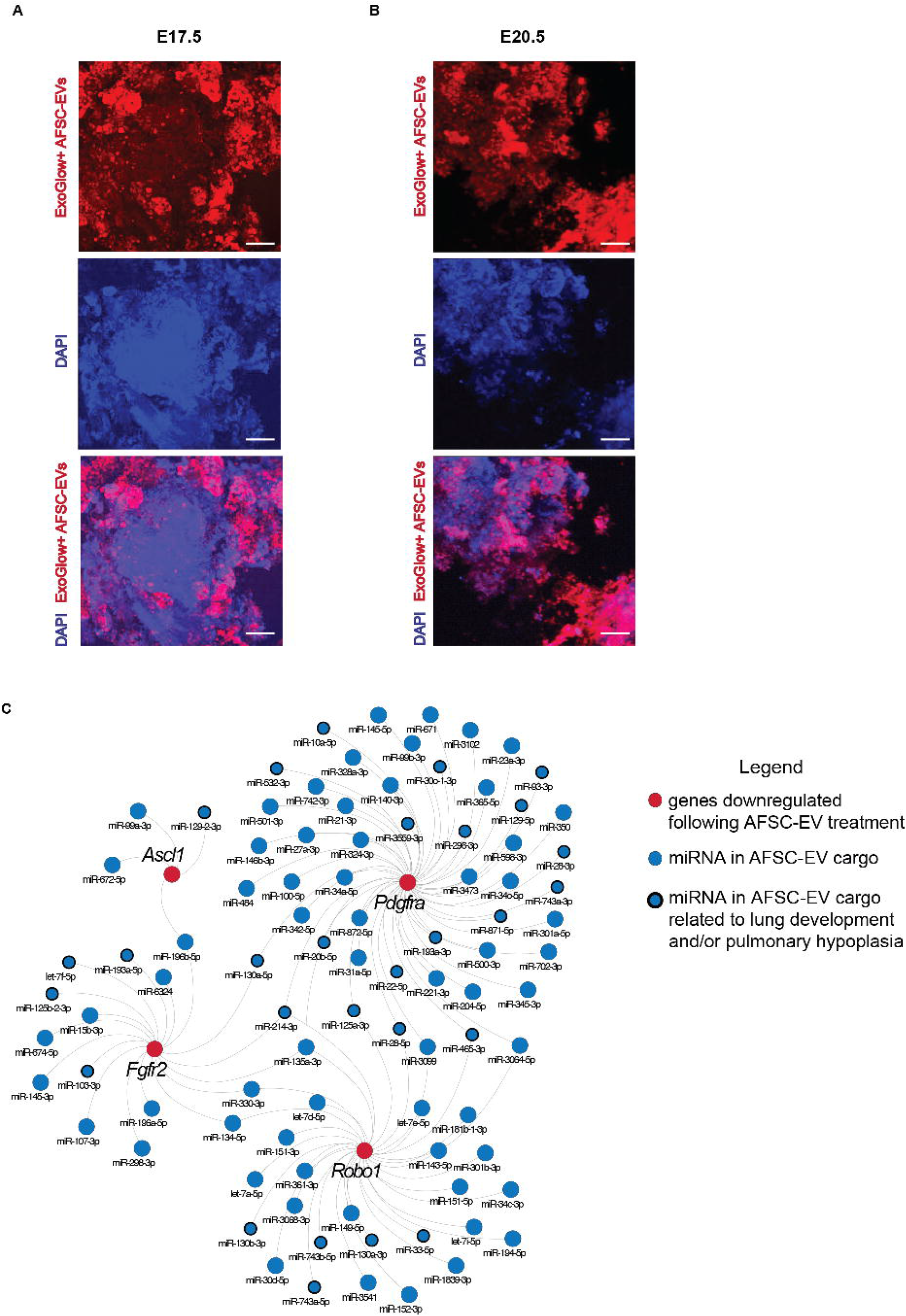
AFSC-EVs modulate fetal lung development and maturation in part through their release of RNA species. Representative images of tracking studies using fluorescently labelled AFSC-EVs (red) in A, E17.5 and B, E20.5 fetal lung explants. DAPI was used as a nuclear stain. C, Network analysis of down-regulated genes responsible for branching morphogenesis and cell differentiation following AFSC-EV treatment (*Robo1, Fgfr2, Ascl1*, and *Pdgfra)* with AFSC-EV cargo miRNA. Scale bar = 50 μm.

## Discussion

In this study, we have shown that AFSC-EVs administered to rat fetal hypoplastic lungs harvested at canalicular and saccular stages of lung development exert a regenerative effect on the pulmonary epithelium and mesenchyme. These effects were through multiple pathways that are known to be involved in lung development [8] and that we herein demonstrated to be targeted by AFSC-EVs. When we investigated the FGF pathway in the nitrofen model, we replicated the observation of other research groups that reported a consistent decrease in FGF10 expression at both canalicular and saccular stages [28–32], and an upregulation of FGFR2 expression at the saccular stage [33]. As we had previously demonstrated in fetal lung explants harvested at the pseudoglandular stage [7], AFSC-EVs administered to explants harvested at both canalicular and saccular stages rescued the expression of both FGF10 and FGFR2 back to control levels. The ROBO/SLIT pathway is also essential for normal lung development, as shown in knock-out studies [34,35], and is downregulated in nitrofen-exposed lungs of fetuses with CDH [36]. In the present study, we confirmed the upregulation of ROBO1 and the corresponding downregulation of ROBO2 and SLIT2 expression in nitrofen-exposed lungs at both timepoints, which were rescued by AFSC-EV treatment to control levels. When we assessed factors involved in sending guidance cues for lung branching, we confirmed that the Ephrin pathway was downregulated at the saccular stage, as previously reported [37], and rescued by AFSC-EV treatment to control levels. Neuropilin is another guidance cue factor that has been reported to support the formation of alveolar ducts and alveoli in rodents [38] and that, to the best of our knowledge, has not been investigated in nitrofen-exposed lungs. The present study shows for the first time that Neuropilin gene expression is downregulated in nitrofen-exposed lungs during the saccular stage and rescued by AFSC-EV administration. Similarly, β-Catenin, which is important for the maintenance of lung epithelial progenitors [39], is herein reported for the first time to have downregulated expression in nitrofen-exposed lungs. However, treatment with AFSC-EVs did not affect β-Catenin gene expression levels. Lastly, when we investigated other factors implicated in lung branching morphogenesis, such as Wnt family members WNT3A and WNT5A, as well as Semaphorin, and Plexin A [8], we found no differences between control and nitrofen-exposed lungs. These results support the notion that some but not all of the pathways involved in branching morphogenesis are dysregulated in human and experimental CDH.

In this study, we confirmed that fetal hypoplastic lungs have impaired epithelial cell differentiation, as we had reported at the pseudoglandular stage [7]. In the distal airways, we found reduced expression of AT1 and AT2 markers at both canalicular and saccular stages of lung development, as previously reported by other groups [25,40–42]. AT1 and AT2 downregulation was rescued by AFSC-EV administration to control levels, in line with our previous report in fetal lungs harvested at the pseudoglandular stage [7]. When we investigated epithelial cell populations in the proximal airways, we observed no differences in the expression of club, basal, and ciliated epithelial cells at the canalicular stage, similar to reports from human and rodent models of pulmonary hypoplasia at the same timepoint [25,43]. At the saccular stage, we observed downregulated expression of club and ciliated epithelial cell markers, as well as upregulated expression of pulmonary neuroendocrine cells in nitrofen-exposed lungs harvested. To the best of our knowledge, no study investigated the expression of basal, club, and ciliated epithelial cells in nitrofen-exposed lungs at the saccular stage. Conversely, we confirmed the upregulation of pulmonary neuroendocrine cells at the saccular stage of lung development, as shown by several groups [44–46]. The increased number of pulmonary neuroendocrine cells in CDH fetal lungs has previously been explained as an adaptive response to impaired lung development, whereby the amines and peptides secreted by these cells are implicated in lung development, repair, and regeneration [44,47]. The dysregulated expression of epithelial cell populations lining the proximal airway were restored to normal levels by AFSC-EV administration at both canalicular and saccular stages.

The mesenchyme plays a critical role in lung development, as it provides support for the fetal lung epithelium [48–50]. In this study, we confirmed that hypoplastic CDH lungs have decreased expression of lipofibroblasts, as previously shown in the nitrofen model [19–21,23] and in the surgical rabbit model of CDH [7]. Lipofibroblasts, a population of lipid-containing fibroblasts that develop during the canalicular stage, play a crucial role during fetal lung development, maintain tissue homeostasis, and respond to cellular injury [48,51]. PDGFRA+ cells are a population of progenitors that mainly give rise to myofibroblasts [27]. This cell population is key for lung development and its conditional inactivation during embryogenesis has been reported to result in alveolar simplification [27]. In our study, we found increased levels of PDGFRA *in vitro* and *ex vivo* at the canalicular stage, as previously reported during the same developmental stage by Dingemann et al [24]. The alteration in markers of lipofibroblasts and PDGFRA+ cells in nitrofen-exposed lungs compared to control could indicate that hypoplastic fetal lungs remain at an immature state, with decreased cell differentiation and increased levels of progenitor cells. This is in line with other studies reporting that the pulmonary mesenchyme is defective in the nitrofen model of pulmonary hypoplasia and may be due to the dysregulated epithelial-mesenchymal tissue interactions [18,42]. The decrease in lipofibroblast markers observed in this study at the canalicular stage persisted also in the saccular stage of lung development. Conversely, we observed that PDGFRA protein expression was decreased in nitrofen-exposed lungs compared to control at the saccular stage. This could be due to simplification of the lung architecture, which is also observed in immature lungs affected by bronchopulmonary dysplasia, where PDGFRA expression has been reported to be downregulated [52–54]. In both canalicular and saccular stages of lung development, AFSC-EV administration to nitrofen-exposed lungs restored markers of lipofibroblasts and PDGFRA+ cells back to control levels. These findings in the mesenchyme were similar to the improvements observed in lung epithelial differentiation. Given the close interaction between epithelium and mesenchyme compartments during lung development, it is difficult to delineate whether EV effects are directed toward one compartment and affects the other indirectly or are affecting both simultaneously.

The EV-tracking experiments conducted in this and our previous study [7] demonstrate that the AFSC-EV cargo uptake is not specific to an area or compartment of the lung. Studies investigating potential cell-type specific transcriptomic changes are currently underway. Nonetheless, in the present study, we were able to find regulators of both lung epithelial and mesenchymal cells. In fact, our analysis of the AFSC-EV miRNA cargo and downregulated genes revealed 28 miRNAs that play a role in restored epithelial and mesenchymal cell differentiation, as well as branching morphogenesis. As an example, miR-20-5p, which belongs to the miR17~92 family, is of particular interest as it controls pulmonary surfactant gene expression in AT2 cells [55]. In addition to this cluster, miR-28-5p was also found in our network to regulate *Robo1* and *Pdgfra*, and has been reported to be decreased in the nitrofen model of CDH [7,56]. Interestingly, 11 miRNAs were identified in our network to regulate more than one of the target genes, indicating that target gene regulation can be coordinated by multiple combinations of miRNAs [57]. Together, these findings support that AFSC-EV mechanism of action in restoring tissue homeostasis is partly ascribed to their RNA cargo transfer.

This study provides further evidence that an AFSC-EV-based therapy holds potential for fetal lung regeneration in babies with pulmonary hypoplasia secondary to CDH. We have herein demonstrated that AFSC-EVs promote branching morphogenesis and rescue epithelial and mesenchymal cell homeostasis at both canalicular and saccular stages of lung development. Towards the translational advancement of this novel therapy, these timepoints are amenable for fetal intervention, especially in fetuses with CDH who are diagnosed prior to these stages. We specifically found that while the changes in differentiation of the distal airways occurred in both the canalicular and saccular stages, the proximal epithelium was rescued only in the saccular stage. However, it is difficult to translate these findings from a rodent model directly to human babies. Fetal interventions in human babies with CDH, such as fetal endoscopic tracheal occlusion, have been reported during both time points [58,59]. Future studies will examine the optimal gestational age to administer the EV-therapy and will address other aspects such as route of delivery and dosage in larger animal models. In fact, it is critical to carefully address these and other translational aspects before clinical application of EV-based therapies in human patients, as previously reported [60,61].

We acknowledge that our study has some limitations. Our findings were obtained using the nitrofen model of CDH, which is based on the administration of an herbicide that targets retinoic acid synthesis [26,62]. Nonetheless, although human pulmonary hypoplasia is not caused by maternal exposure to herbicides, nitrofen administration causes a degree of lung underdevelopment analogous to that of human fetuses with pulmonary hypoplasia [26,62]. Moreover, retinoic acid is known to play a role in fetal lung development and CDH babies have low intracellular retinoic acid levels in the lungs and low retinol and retinol-binding protein levels in the cord blood [63,64]. This model has been used for over three decades and the multiple pathways that are dysregulated in nitrofen-exposed hypoplastic lungs are similar to those reported in human studies [26]. Nevertheless, findings from our study serve mainly as a proof-of-principle that AFSC-EVs rescue lung development in CDH hypoplastic lungs. Moreover, we recognize that our study addressed the epithelial and mesenchymal compartments of the lung, for what is currently known. While there is a vast literature on epithelial differentiation in normal and hypoplastic lungs, relatively little is known about lung mesenchymal cells and their impairment in pulmonary hypoplasia. This is in line with a recent report by Tsukui et al, who highlighted that the diversity of lung fibroblasts has not been well characterized in normal or diseased lungs [50]. Nonetheless, we are encouraged by the observation that AFSC-EVs are able to promote differentiation towards lipofibroblast in canalicular and saccular stages in rats, as well as during alveologenesis in fetal rabbits with CDH[7].

In conclusion, this study confirms that administration of AFSC-EVs to hypoplastic fetal lungs rescues branching morphogenesis and promotes differentiation of epithelial and mesenchymal lung cells during the canalicular and saccular stages of lung development. Moreover, we provide further evidence that the RNA cargo could be responsible for the AFSC-EV mechanism of action towards lung regeneration. Taken together, these findings demonstrate that AFSC-EV treatment may represent an exciting new opportunity for antenatal intervention for CDH babies at clinically relevant timepoints.

## Acknowledgements

The authors would like to thank Rachel Bercovitch for technical assistance and Paolo De Coppi for providing AFSCs in kind. We are indebted to the Imaging Facility and the Lab Animal Services at the Hospital for Sick Children, Toronto. The graphical abstract was designed using BioRender.com.

