## Supplemental Figure Legends for "Amniotic fluid stem cell extracellular vesicles promote fetal lung branching and cell differentiation in experimental congenital diaphragmatic hernia"

**Supplemental Figure and Video legends**

**Supplemental Figure 1. Factors involved in branching morphogenesis not altered in hypoplastic fetal rat lungs.** Gene expression levels (RT-qPCR) of factors implicated in branching morphogenesis which were not differentially expressed by nitrofen nor affected by AFSC-EV administration at the A, canalicular and B, saccular stages of lung development. Data were analyzed using one-way ANOVA using Tukey’s multiple comparison test. Data are presented at mean ± SD (n= at least 3 biological replicates per group).

**Supplemental Figure 2.** **Markers for distal fetal lung epithelium are not altered in hypoplastic fetal rat lungs at the canalicular stage of development.** Gene expression levels of markers of club, basal, ciliated epithelial, and neuroendocrine cells in fetal lung explants at the A, gene (RT-qPCR) and B, protein (Western blot) level at the canalicular stage. ACTB was used as a loading control. Data were analyzed using one-way ANOVA using Tukey’s multiple comparison test. Data are presented at mean ± SD (n= at least 3 biological replicates per group).

**Supplemental online Video 1.** Representative video of EV tracking studies using fluorescently labelled AFSC-EVs (red) at the saccular stage of lung development. DAPI was used as a nuclear stain.
