## Supplementary figures and images for "Amniotic fluid stem cell extracellular vesicles promote fetal lung branching and cell differentiation in experimental congenital diaphragmatic hernia"

### Supplemental Figure 1

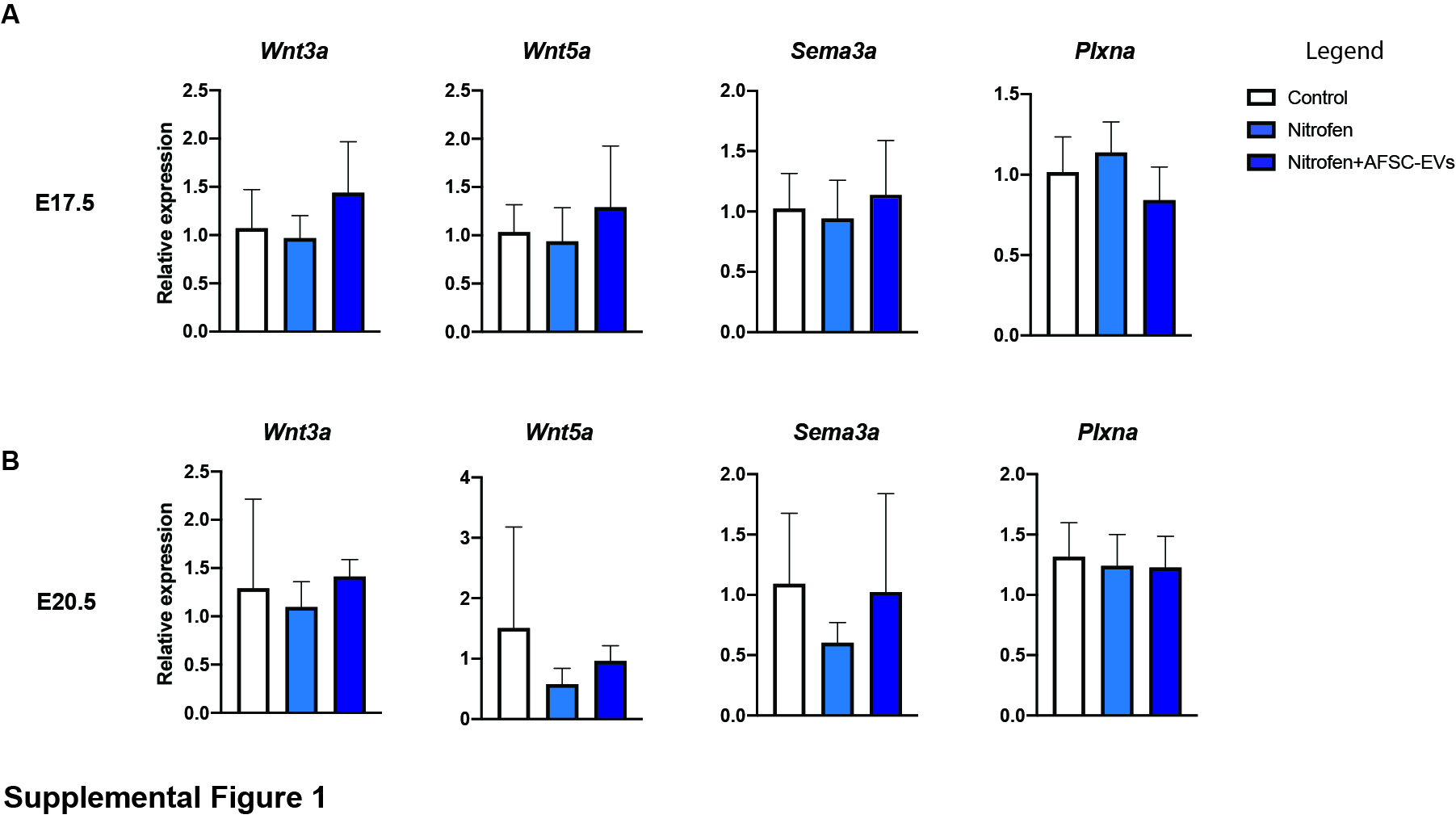

### Supplemental Figure 2

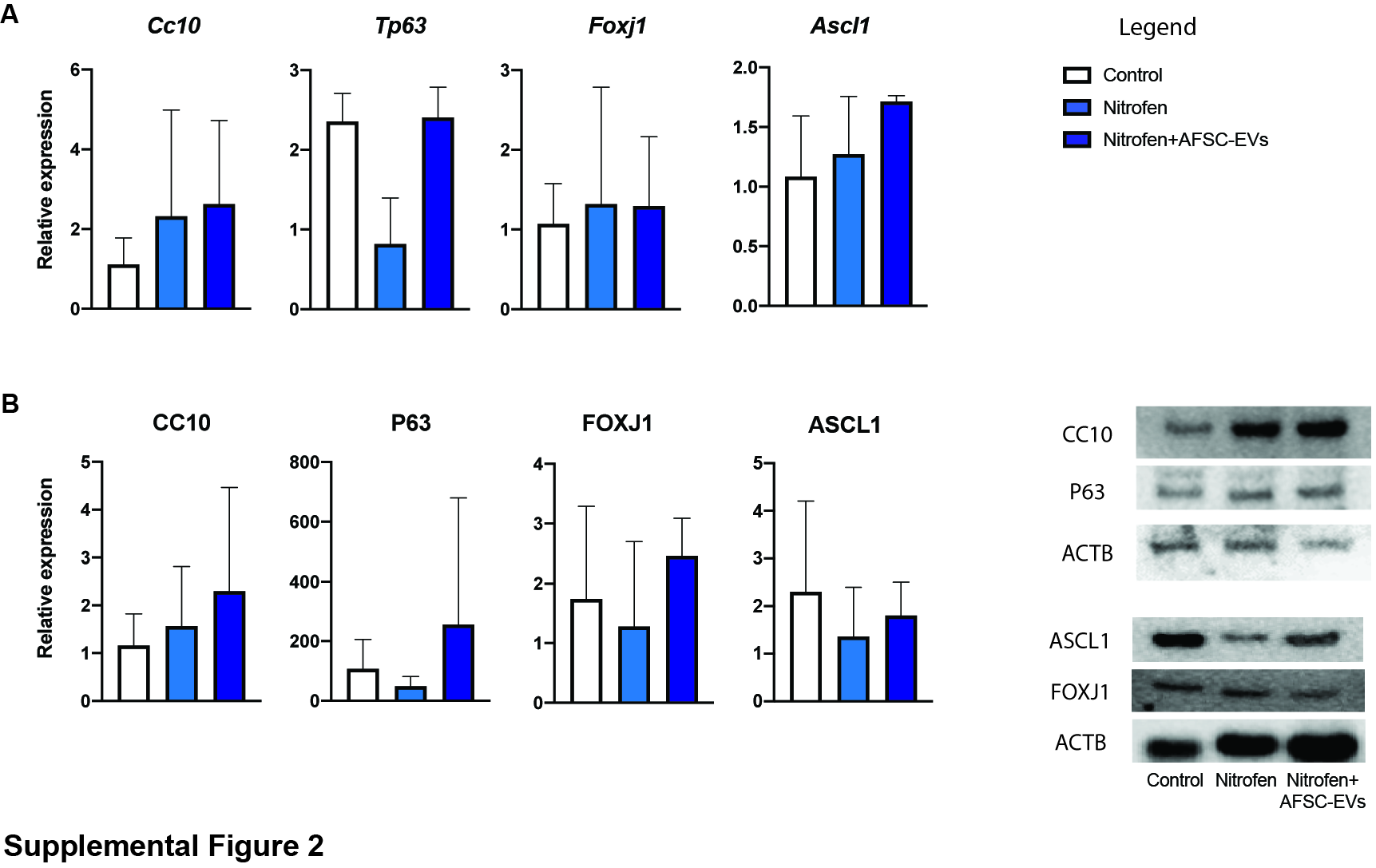
